# Root hair lifespan is antagonistically controlled by autophagy and programmed cell death

**DOI:** 10.1101/2025.03.18.643910

**Authors:** Qiangnan Feng, Shihao Zhu, Xinchao Wang, Yujie Liu, Jierui Zhao, Yasin Dagdas, Moritz K. Nowack

## Abstract

Root hairs are tubular tip-growing extensions of root epidermal cells that enhance root surface area for water and nutrient uptake. While mechanisms governing root hair fate, polarity, and tip growth are well understood, the regulation of root hair longevity remains largely unknown. Here, we show that root hair cells employ high levels of autophagy to promote their lifespan. Loss-of-function mutations in the autophagy regulators ATG2, ATG5, or ATG7 induce a premature, cell-autonomous cell death program. This cell death is activated via a gene regulatory network downstream of the NAC transcription factors ANAC046 and ANAC087. Our findings uncover an antagonistic relationship between autophagy and developmentally controlled cell death in root hair lifespan regulation, with potential implications for optimizing plant nutrient and water uptake in crop breeding.

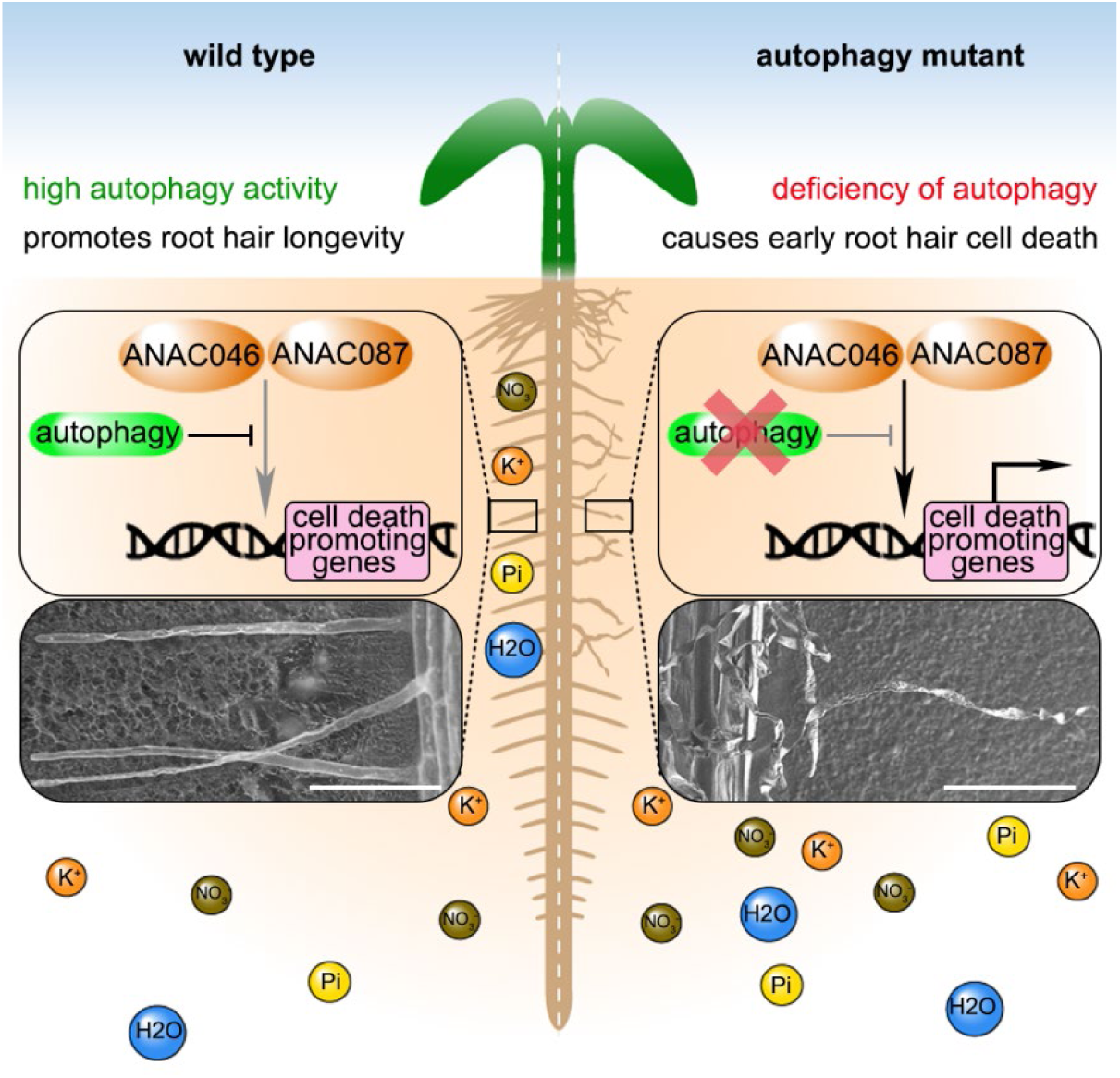

## Introduction

Root hairs are important for plant growth and development, since they dramatically increase the root surface area available for water and mineral uptake, improve root anchoring in the soil, and facilitate signaling with the soil microbiome (Grierson et al., 2014). Root hair development occurs in four phases: cell fate specification, initiation of outgrowth in the basal cell region, tip growth maintenance, and growth arrest at maturity (Gilroy and Jones, 2000). The patterning of epidermal cells in root hair and non-root hair cells, and subsequent cellular differentiation is regulated by extensive gene regulatory networks (Vissenberg et al., 2020). Initiation and maintenance of root hair tip growth depends on additional transcription factors, cell wall-modifying enzymes, gradients of calcium and reactive oxygen species, and targeting of active ROPs to specific plasma membrane domains (Honkanen and Dolan, 2016; Zhang et al., 2022; Li et al., 2023). After having reached a certain length, root hairs enter the maturation stage, characterized by vacuolization of the apical root hair region and tip-growth arrest (Grierson et al., 2014). Subsequently, protoplast shrinkage and DNA fragmentation occur, hinting at a controlled degeneration process (Shishkova and Dubrovsky, 2005; Li et al., 2016; Tan et al., 2016). However, the mechanisms and pathways that control root hair life span remain largely unknown.

Both autophagy and programmed cell death have been established as pathways determining cellular life span and organ senescence in animals and plants (Woo et al., 2019; Cassidy and Narita, 2022). Regulated cell death, or programmed cell death (PCD) is a highly organized process occurring in the context of regular development (dPCD), and as part of the reaction to environmental stresses or insults (ePCD) (Daneva et al., 2016). While the regulation of numerous PCD subroutines have been established in animal systems (Galluzzi et al., 2018), the molecular control of PCD in plants appears not to be conserved and is much less understood (Kacprzyk et al., 2024). Plant dPCD is controlled by gene regulatory networks which control the expression of a suite of specific dPCD signature genes prior to PCD execution (Olvera-Carrillo et al., 2015). NAC transcription factors play key roles in controlling the timely onset of dPCD in several developmental contexts. Some coordinate tissue differentiation and dPCD preparation in a highly cell-type specific manner, e.g. SMB in the root cap (Fendrych et al., 2014) or VNDs in the xylem (Kubo et al., 2005; Ohashi-Ito et al., 2010). Others NAC factors appear to act further downstream in the dPCD regulatory network and are commonly upregulated in several dPCD contexts independent of the tissue type, like ANAC046 and ANAC087 that have been implicated in dPCD processes in the xylem, the root cap, stigmatic papilla cells, the endosperm, and leaves (Kim et al., 2014; Oda-Yamamizo et al., 2016; Gao et al., 2018; Huysmans et al., 2018; Doll et al., 2023).

In contrast to dPCD pathways, autophagy is highly conserved in eukaryotes, and regulated by a suite of established autophagy genes (ATGs). Autophagy acts as a quality control pathway, mediating the degradation of cellular components by targeting them to the lysosomes or vacuoles (Gross et al., 2025). In plants experiencing optimal growth conditions, autophagy operates at basal levels to maintain cellular homeostasis, but starvation and environmental stresses can upregulate autophagic activity to promote plant survival (Agbemafle et al., 2023). In contrast to these pro-survival functions, autophagy can promote the transition to cell death, for instance under nutrient starvation conditions in cell cultures (Teper-Bamnolker et al., 2021). In plant immunity, autophagy has been assigned pro-survival as well as pro-death roles, depending on context (Liu et al., 2005; Hofius et al., 2009). In the Arabidopsis root cap, autophagy has cell-type specific roles: It promotes cell sloughing as well as cell death and corpse clearance in root cap columella cells, but is dispensable for dPCD execution and corpse clearance in lateral root cap cells (Feng et al., 2022; Goh et al., 2022).

Here, we report that defects in the canonical autophagy pathway of Arabidopsis causes the premature onset of dPCD signature gene expression specifically in root hairs. In line with this observation, we find that root hair longevity in different *atg* mutants is reduced by precocious activation of a root-hair specific dPCD process. Cell type-specific ATG knockout and transgene complementation confirmed the cell-autonomous pro-survival role of autophagy in root hairs. Interestingly, root hair cell death depends on ANAC046 and ANAC087, linking the pro-survival role of autophagy in root hairs to the suppression of canonical dPCD pathways.

## Results

### Root hair cells display intrinsically elevated autophagic activity

Investigating root cell-type specific differences in autophagy processes, we imaged roots of *Arabidopsis thaliana* (Arabidopsis) wild-type and *atg* mutants expressing the autophagy reporter *YFP-ATG8A*. In the wild-type, we detected YFP-ATG8A both freely cytosolic and in a high number of punctate foci representing autophagosomes (APGs) in different root hair developmental stages (Figure 1A). By contrast, in *atg7-2* and *atg5-1* mutants, we detected a predominantly cytosolic YFP signal and significantly less APGs in mature root hair cells (Figure 1B-D), suggesting an upregulation of canonical autophagy in this cell type. Using Concanamycin A (ConA) as a drug to allow quantification of autophagic delivery to the central vacuole, we counted significantly more YFP-ATG8A-positive autophagic bodies in vacuoles of root hair cells than non-root hair cells, indicating intense autophagy activity also in young root hair cells (Figure 1E-F), suggesting that autophagy is highly activated throughout root hair development.

**Figure 1.**
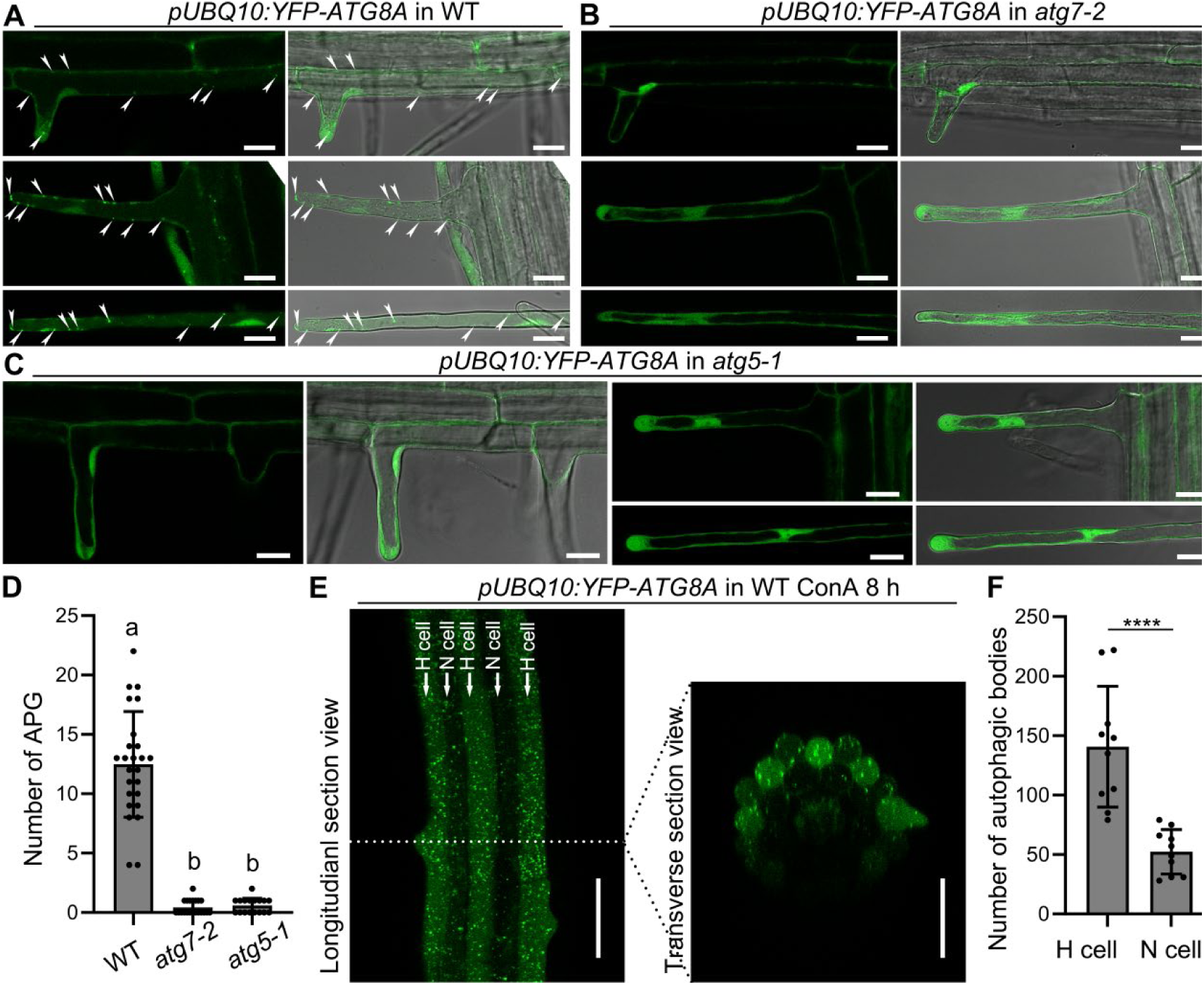
Autophagy activation is increased in root hair cells. A-C, Representative confocal images of root hair cell from 5 days after germination (DAG) expressing *pUBQ10:YFP-ATG8A* in wild-type (WT) (A), *atg7-2* (B) and *atg5-1* (C). White arrows indicate autophagosomes (APGs). D, The quantification of APG in mature root hair cells. Results are means ± SD. 17-25 root hair cells from 5 roots of each genotype were analyzed. Means with different letters are significantly different (one-way ANOVA, Tukey’s multiple comparisons test, P < 0.05). E, Representative confocal images of roots from seedlings at 5 DAG expressing *pUBQ10:YFP-ATG8A* in WT, *atg7-2* and *atg5-1* mutant treated with 1 μM ConA for 8 h. Non-root hair cell (atrichoblast) and root hair cell (trichoblast) were shown as N cell and H cell, respectively. F, The quantification of autophagic bodies in H cells treated with 1 μM ConA for 8 h as shown in (E). Results are means ± SD. N = 10 cells from 5 roots. **** indicates a significant difference (*t* test, P < 0.0001). Bars = 20 μm for A-C, 50 μm for E.

### Autophagy deficiency activates a premature dPCD process in root hair cells

To investigate functions of autophagy in root hairs, we investigated root hairs of wild-type and *atg* mutants. While *atg* mutant root hair initiation and tip growth was indistinguishable from the wild type (Supplemental Figure 1A-D), we found collapsed root hairs in all but the most distal parts of 7-day old seedling roots (Figure 2A). Next, we generated transgenic lines expressing a ubiquitously expressed *pUBQ10:ATG5-mCherry* complementation construct in the *atg5-1* mutant background. Three independent lines showed a complete restoration of the root hair phenotype (Figure 2B), suggesting that autophagy is crucial determinant of root hair cell longevity in Arabidopsis.

**Figure 2.**
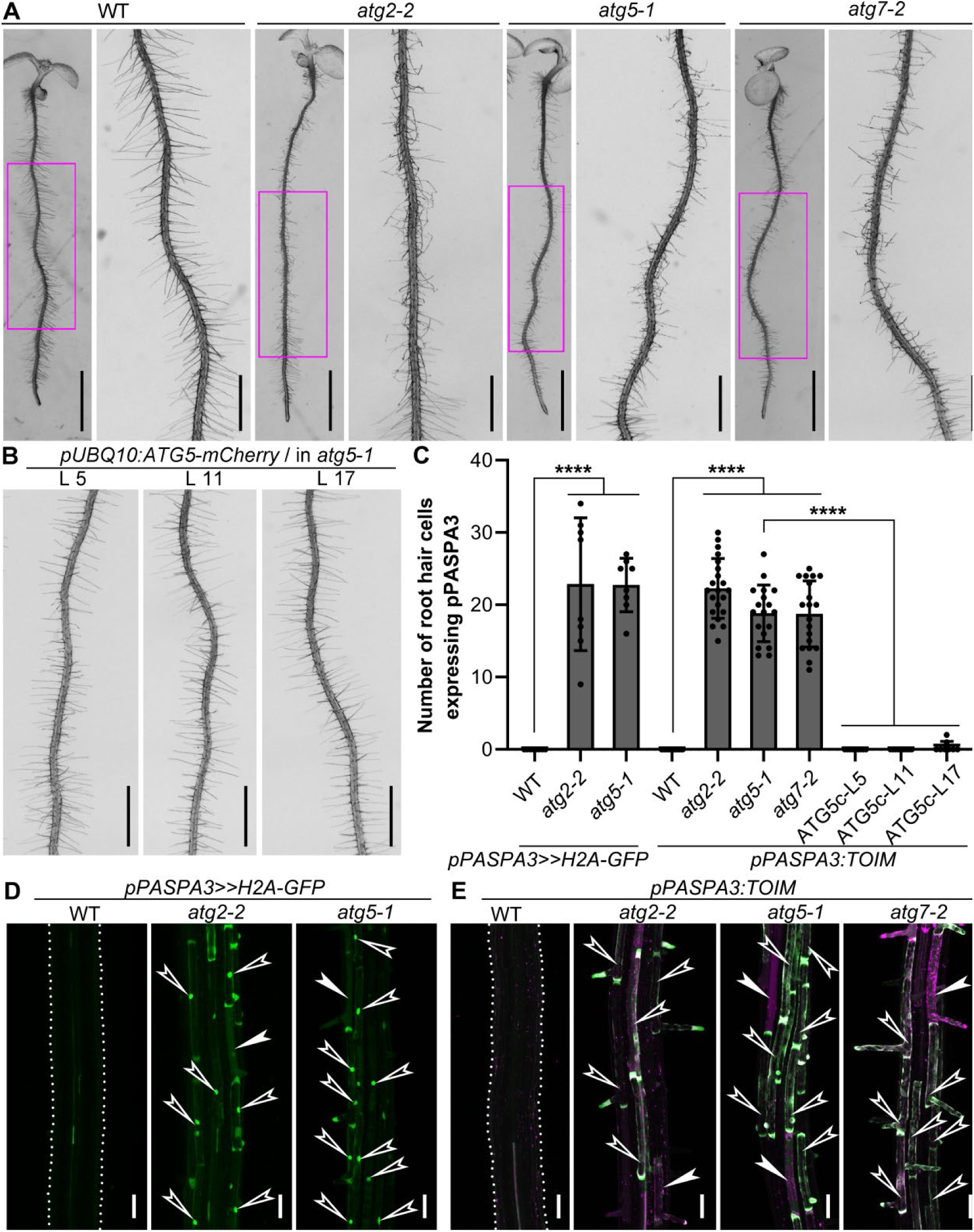
The loss function of autophagy promotes precocious expression of PASPA3 and dPCD onset in root hairs. A, Representative primary roots from 7-day-old seedlings: WT, *atg2-2*, *atg5-1* and *atg7-2*. The right panel of each genotype shows a zoomed image of a part region of the left panel outlined in magenta. B, Representative primary roots from 7-day-old seedlings: three independent lines of *pUBQ10:ATG5-mCherry* in *atg5-1* background. C, The quantification of root hair cells expressing PASPA3 in WT, *atg* mutants and complement lines (ATG5c). The name of lines *pUBQ10:ATG5-mCherry* in *atg5-1* expressing *pPASA3:TOIM* was written as ATG5c. In total, 8-23 roots were analyzed for each genotype. **** indicates a significant difference (*t* test, P < 0.0001). D-E, Representative confocal images of roots from 5-day-old WT and *atg* mutants expressing *pPASPA3>>H2A-GFP* (D) or *pPASPA3:TOIM* (E). Arrowheads outlined in white indicate that H2A-GFP (D) signals or TOIM (E) were observed in root hair cells of *atg* mutants. White arrowheads indicate that nuclear leakage (D) or vacuole collapse (E) of *atg* mutants. White dotted lines point the profile of root in WT. Bars = 5 mm for A (left panel of each genotype), 1 mm for A (right panel of each genotype) and B, 50 μm for D and E. See also Figure S1 and S2; Movie S1 and S2.

To test if root hair collapse displayed dPCD hallmarks, we introgressed promoter-reporters of the dPCD signature genes *PASPA3 and RNS3* (Fendrych et al., 2014; Olvera-Carrillo et al., 2015) into *atg* mutants. While PASPA3 and RNS3 expression in the wild type is restricted to root cap and xylem cells preparing for dPCD (Fendrych et al., 2014; Olvera-Carrillo et al., 2015), *atg* mutants displayed ectopic PASPA3 and RNS3 expression in young root hair cells (Figure 2D-E, Supplemental Figure 1E-F). Notably, the ectopic PASPA3 expression is not present in ATG5-mCherry complementation lines (Figure 2C). In line with reporter expression, we could observe cellular hallmarks of dPCD (Fendrych et al., 2014; Wang et al., 2024), including mitochondrial disintegration, nuclear envelope breakdown, and vacuolar collapse, in *atg* mutant root hairs (Figure 2D-E, Supplemental Figure 1G-H). As root hairs can be easily damaged by microscopy mounting, we followed root hair viability and death in seedlings grown in chambers that allow minimally invasive imaging of plants. In line with our previous results, *atg5-1* mutant root hairs were dead, while wild-type root hairs in the same root region were alive (Supplemental Figure 2A). To address the question whether dPCD in *atg* mutant root hairs occurs ectopically or rather prematurely, we imaged root hairs of 14-day old plants during different stages of secondary growth and periderm development (Wunderling et al., 2018). We found that root hair cells expressed the PASPA3 reporter and subsequently collapsed in the most mature root region just below the hypocotyl (Supplemental Figure 2B, C and F). In *atg2-2* mutants, by contrast, root hairs expressed PASPA3 and subsequently collapsed in a much earlier stage (Supplemental Figure 2D, E and F), suggesting that autophagy deficiency severely reduces root hair life span by early activation of senescence-related dPCD.

### Autophagy promotes root hair longevity in a cell-autonomous fashion

Seeing that autophagy is a systemically active process in plants, we next addressed the question if premature root hair cell dPCD was caused by lack of autophagy specifically in root hairs, or else a more indirect, non-cell autonomous effect of systemic autophagy deficiency. To this end, we generated root-hair specific complementation lines, expressing ATG5-mCherry under the root hair-specific promoter of *ARABIDOPSIS THALIANA EXPANSIN A7* (pE7) (Cho and Cosgrove, 2002; Won et al., 2009). As autophagy has been implicated in xylem maturation (Kwon et al., 2010) and in the restriction of cell death to specific xylem cell types (Escamez et al., 2016; Escamez et al., 2019), we used the xylem-specific promoter IRREGULAR XYLEM1/CELLULOSE SYNTHASE8 (pIRX1) (Turner and Somerville, 1997; Taylor et al., 2000) controlling ATG5-mCherry to test if root hair cell death was dependent on xylem autophagy (Figure 3A). We established three independent lines of *pE7:ATG5-mCherry* that showed a complete rescue of the *atg5-1* phenotype, both the early expression of PASPA3 (Figure 3B) and root hair degeneration (Figure 3C). By contrast, we could not find any line of *pIRX1:ATG5-mCherry* that showed complementation (Figure 3B and D).

**Figure 3.**
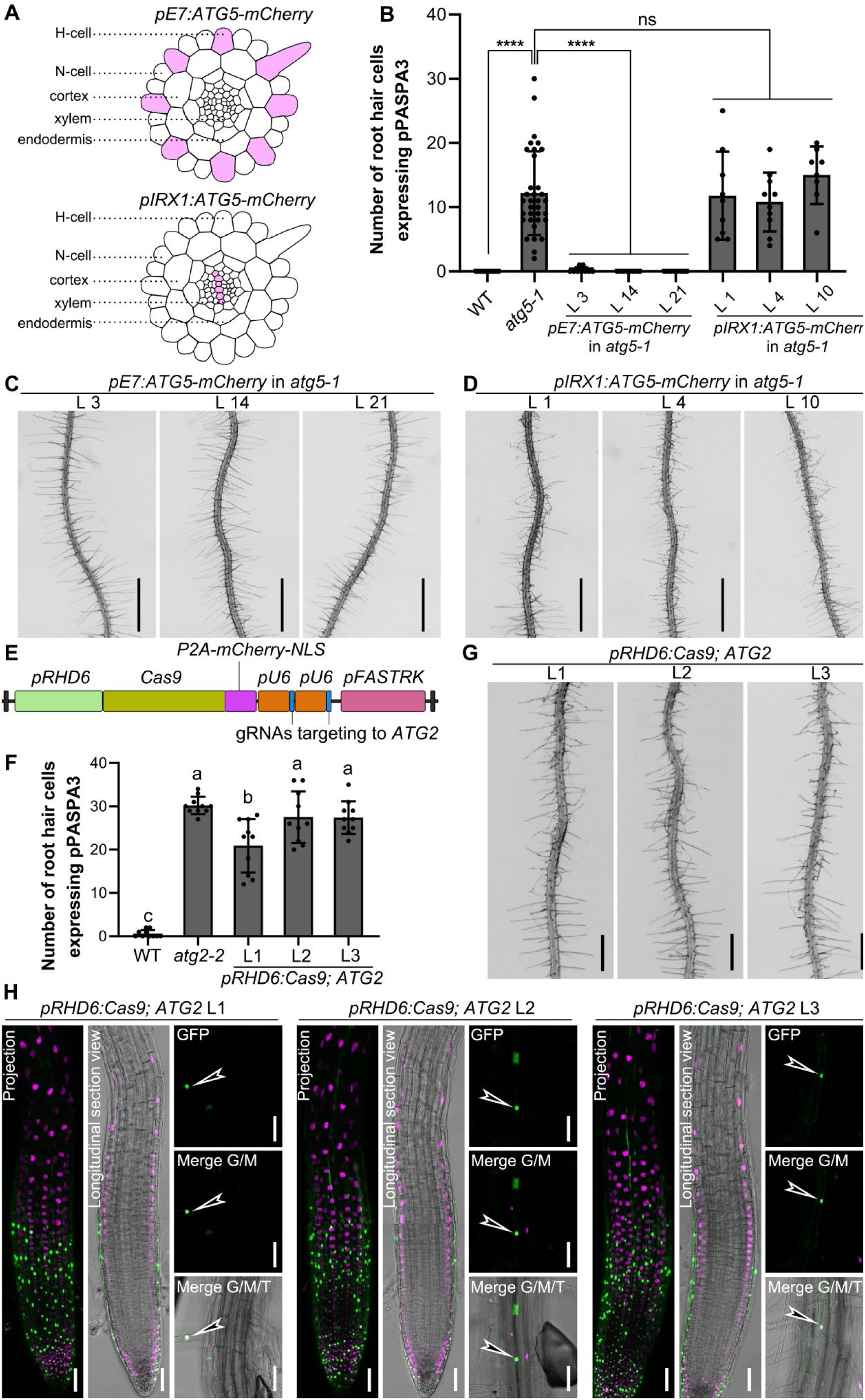
Root-hair-specific complementation and loss of function shows root hair inherent functions of autophagy. A, Schematic representation of the cross-section of Arabidopsis root. The expression pattern of *pE7:ATG5-mCherry and pIRX1:ATG5-mCherry* is shown as pink in the upper panel and lower panel, respectively. B, The quantification of root hair cell expressing *PASPA3*. Results shown are means ± SD. In total, 8-34 roots were analyzed for each genotype. **** indicates a significant difference (*t* test, P < 0.0001). ns indicates not significant (*t* test, P > 0.05). C-D, Representative primary roots from 7-day-old seedlings: three independent lines of *pE7:ATG5-mCherry* (C) and *pIRX1:ATG5-mCherry* (D) in *atg5-1* background. E, Schematic representation of the vector: *pRHD6:Cas9;ATG2.* The Cas9-P2A-mCherry is driven by root hair specific promoter pRHD6, and two gRNAs targeting to *ATG2* are inserted. F, The quantification of root hair cell expressing *PASPA3*. Results shown are means ± SD. In total, 10 roots were analyzed for each genotype. Means with different letters are significantly different (one-way ANOVA, Tukey’s multiple comparisons test, P < 0.05). G, Representative primary roots from 7-day-old seedlings: three independent lines of *pRHD6:Cas9; ATG2*. H, Representative confocal images of three independent lines of *pRHD6:Cas9;ATG2* in *pPASPA3>>H2A-GFP* background. Arrowheads outlined in white indicate the precocious expression of *PASPA3* in root hair cells. Bars = 1 mm for C and D, 500 μm for G, 50 μm for H. See also Figure S3.

Next, we used a tissue-specific knockout approach, CRISPR-TSKO (Decaestecker et al., 2019; Bollier et al., 2021), expressing *Cas9* and a polycistronically attached *P2A-mCherry-NLS* sequence under the pE7 promoter. Combined with previously reported guide RNAs (gRNAs) targeting *ATG2* (Feng et al., 2022), we generated *pE7:Cas9;ATG2* (Supplemental Figure 3A). The construct was transformed into wild-type plants carrying *pPASAP3>>H2A-GFP* (Fendrych et al., 2014), and two independent lines with high expression levels of mCherry-NLS in root hair cells were established (Supplemental Figure 3B). While these lines showed a significant rise of *pPASPA3>>H2A-GFP* activity in root hairs, they did only partially phenocopy the premature root hair collapse of *atg2-2* mutants (Supplemental Figure 3C-D). To test if this is due to the late activation of *pE7* in the course of root hair development, we repeated the experiment with Cas9 driven by the promoter of *ROOT HAIR DEVELOPMENT6* (*pRHD6*) (Figure 3E), which is activated already in the immature root hair cell files in the root meristem (Menand et al., 2007). Three independent lines of *pRHD6:Cas9;ATG2* showed not only early pPASPA3 reporter expression (Figure 3F and 3H), but also a root hair degradation phenotype similar to the *atg2-2* mutant (Figure 3G). Taken together, these results demonstrate a cell-autonomous function of autophagy in promoting root hair longevity by suppressing senescence-induced dPCD.

### Autophagy-deficient root hair degeneration is SA-, JA- and ethylene-independent

The early leaf senescence phenotypes seen in *atg* mutants depends on salicylic acid (SA) signaling (Yoshimoto et al., 2009). To investigate possible analogies between leaf and root hair senescence, we crossed the SA-biosynthesis mutant *sid2* (Nawrath and Métraux, 1999) with the *atg2-2* mutant expressing a *pPASPA3* reporter. Similar to *atg2-2*, the *atg2-2 sid2* double mutant shows early PASPA3 expression and root hair degeneration (Supplemental Figure 3E-G). This was confirmed by the *atg5-1 sid2* double mutant (Supplemental Figure 3H), showing that the *atg* mutant root hair dPCD phenotype does not depend on SA signaling.

Also jasmonic acid (JA) and ethylene have been implicated in senescence modulation (Liao et al., 2022), prompting us to test their involvement in premature root hair degeneration by generating double mutants of *atg2-2* and *atg5-1* with *coi1* (Huang et al., 2014) and *ein2* (Alonso et al., 1999), respectively. All *coi1* and *ein2* double mutant combinations retained the expression of PASPA3 and early cell death phenotype (Supplemental Figure 3), suggesting that neither SA-, nor JA- or ethylene-signaling pathways are responsible for the premature root hair cell death in autophagy mutants.

### Autophagy-deficient root hair cell death depends on canonical dPCD regulators

To test the involvement of established dPCD-regulating transcription factors in autophagy-deficient root hair cells, we interrogated their expression patterns using published single-cell RNA-sequencing (scRNA-seq) datasets (Denyer et al., 2019; Wendrich et al., 2020). We found that both *ANAC046* and *ANAC087* are expressed in sc-RNAseq clusters corresponding to root hair cells (Supplemental Figure 4A-C). Investigating the published promoter-reporter lines *pANAC046:NLS-tdTOMATO* and *pANAC046:NLS-tdTOMATO* (Huysmans et al., 2018), we confirmed the root hair cell expression of both transcription factors (Supplemental Figure 4D). Next, we investigated 14-day old roots of wild-type and *anac046 anac087* double mutant plants. Compared with wild type, we found significantly less collapsed root hair cells in the hypocotyl-proximal region of the *anac046 anac087* mutant, suggesting the root hairs of *anac046 anac087* remained viable longer than the wild type ones (Supplemental Figure 4E-F). In addition, we generated *atg2-2 anac046 anac087* triple mutants expressing the PASPA3 reporter (*tri-1*). In parallel, we generated two novel null mutant alleles of *ATG2* (*atg2-8* and *atg2-9*) using published gRNAs (Feng et al., 2022) directly in the *anac046 anac087* double mutant background (*tri-2*, *atg2-8 anac046 anac087*; *tri-3, atg2-9 anac046 anac087*) (Figure 4A, Supplemental Figure 5A). Interestingly, we observed that loss of *ANAC046* and *ANAC087* function was sufficient to suppress the premature root hair cell death phenotype of *atg2* mutants (Figure 4B), but not the early leaf senescence of *atg* mutants (Supplemental Figure 5B-C). By digital-droplet PCR, we demonstrated dPCD signature genes were down-regulated to wild-type levels in these triple mutants when compared with *atg2-2* mutant (Figure 4C), suggesting that NAC transcription factors are indispensable for autophagy-mediated root hair dPCD. By quantifying the number of root hair cells expressing pPASPA3 in *tri-1*, we found that *anac046 anac087* rescues the precocious expression of pPASPA3 of *atg2-2*, confirming the digital-droplet PCR results (Figure 4D).

**Figure 4.**
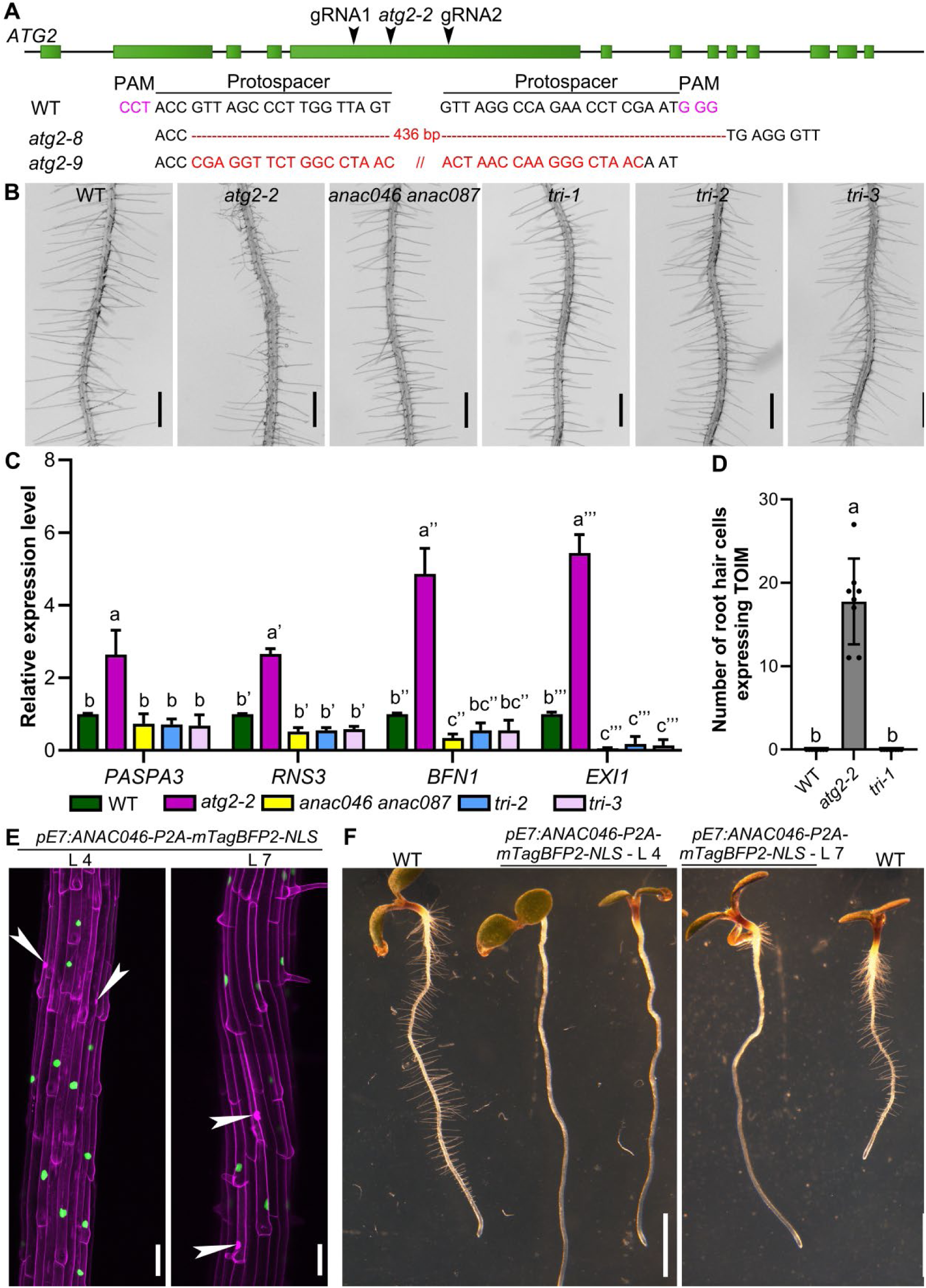
The premature dPCD of root hairs in *atg* mutants dependents on established dPCD gene regulatory networks. A, Schematic illustration of genomic regions of *ATG2*. The *atg2-2* mutation site and Cas9-targeting site are indicated by arrowheads on the genomic loci. Green boxes indicate exon of ATG2. PAM sequences and protospacer sequences are indicated by magenta and black letters, respectively. B, Representative primary roots from 7-day-old seedlings: WT, *atg2-2, anac046 anac087* and three triple mutant lines of *atg2-2 anac046 anac087* (*tri-1*), *atg2-8 anac046 anac087* (*tri-2*) and *atg2-9 anac046 anac087* (*tri-3*). C, Relative expression of PCD-associated genes (*PASPA3*, *RNS3*, *BFN1*, and *EXI1*) in roots of WT, *atg2-2, anac046 anac087, tri-2* and *tri-3* by digital-droplet PCR (ddPCR) analysis. Results shown are means ± SD. Three independent biological replicates (two-three technical repeats each) were performed. Means with different letters are significantly different for each gene (two-way ANOVA, Tukey’s multiple comparisons test, P < 0.005). D, The quantification of root hair cell expressing *PASPA3*. Results shown are means ± SD. In total, 8 roots were analyzed for each genotype. Means with different letters are significantly different (one-way ANOVA, Tukey’s multiple comparisons test, P < 0.05). E, Representative confocal images of roots from 5-day-old seedlings expressing *pE7:ANAC046-P2A-mTagBFP2-NLS* (shown as green) pulse labeled with PI (shown as magenta). White arrowheads point the PI entry. F, Representative images of 5-day-old seedlings: WT and two lines of *pE7:ANAC046-P2A-mTagBFP2-NLS*. Bars = 500 μm for B, 50 μm for E, 2 mm for F. See also Figure S4 and S5.

Finally, we misexpressed ANAC046 under the pE7 promoter, and established two independent lines with root hair cell death in the early stage of root hair formation (Figure 4E). Visually, these lines appear root-hair less (Figure 4F), and show that ANAC046 expression is sufficient to cause root hair cell death.

In sum, these results demonstrate the premature dPCD of root hairs in *atg* mutants dependents on canonical dPCD gene regulatory networks that are shared with established dPCD processes as they occur in the xylem, the root cap, or the endosperm.

## Discussion

Root hairs as tubular extensions of epidermal cells can represent a substantial portion of the root system surface area (Bates and Lynch, 1996). By increasing the interface area between plant and soil with minimal biomass investment, root hairs crucially contribute to water and mineral acquisition in land plants (Duddek et al., 2023). Both the length and number of root hairs have been implicated in optimizing their function (Cai and Ahmed, 2022). Here, we investigated an additional dimension of root hair performance: their functional longevity. Prompted by the discovery of elevated autophagic activity specifically in root hair cells, we found that autophagy maintains root hair viability by delaying a senescence-induced developmental cell death program.

Root hair development from patterning of fate determination to initiation, elongation by tip growth, and maturation has been intensively studied (Cui et al., 2018). However, little is known on developmentally or environmentally controlled senescence of root hairs, and root age-related processes as a whole has received little attention (Tunc and von Wirén, 2025). In cotton, root hair life span has been determined to last about 3 weeks, depending on the soil moisture content (Xiao et al., 2020). Root hair senescence in Arabidopsis has been described to occur after 2-3 weeks, showing hallmarks of programmed cell death including protoplast retraction from the cell wall and nuclear DNA fragmentation (Li et al., 2016; Tan et al., 2016). It has been shown that root hairs of rice (Oryza sativa) and maize (Zea mays) shrink or collapse in response to reduced soil water contents (Keyes et al., 2017; Duddek et al., 2022). In Arabidopsis, senescing root hairs appear to twist and shrink, reducing less water-holding capacity in comparison with turgid root hairs (Li et al., 2016; Choi and Cho, 2019). Hence, research into the factors determining root hair longevity can reveal new angles towards optimizing plant nutrient use and drought resilience of plants.

Despite having been proposed as a model for PCD research upon stress treatments (Hogg et al., 2011), genes controlling root hair longevity and senescence-induced cell death remained unknown. Our results indicate that autophagy is a key factor to determine root hair lifespan and senescence in Arabidopsis, adding a central aspect to the physiological and developmental roles of autophagy.

Mutants of core autophagy components in Arabidopsis and maize show premature leaf senescence, but otherwise develop normally under optimal growth conditions (Marshall and Vierstra, 2018). The involvement of autophagy in cellular pro-life and pro-death processes has been discussed controversially (Üstün et al., 2017). In particular, it has become clear that autophagy is not required for major dPCD events such as xylem biogenesis or lateral root cap turnover, despite the upregulation of autophagic activities in these contexts (Courtois-Moreau et al., 2009; Feng et al., 2022). On the contrary, autophagy in leaf senescence has to maintain cellular viability longevity through nutrient recycling, stalling cell death until nutrient mobilization has been completed (Masclaux-Daubresse et al., 2017).

The NAC transcription factors ANAC046 and ANAC087 have been implicated in leaf senescence control (Vargas-Hernández et al., 2022), controlling chlorophyll breakdown by activating known senescence promoters such as NON-YELLOW COLORING1 or STAY-GREEN1 (SGR1) and SGR2 (Oda-Yamamizo et al., 2016). In a differentiation-induced dPCD context such as the root cap, ANAC046 controls expression of dPCD signature genes BIFUNCTIONAL NUCLEASE1, EXITUS1, and RIBONUCLEASE3 (Olvera-Carrillo et al., 2015; Huysmans et al., 2018).

Similar dPCD signatures have been found in root cells during the onset of secondary growth. Interestingly, dPCD signature genes are expressed in root endodermis cells, but have not been found in epidermal root hair or non-root hair cells (Wunderling et al., 2018). As the onset of secondary growth precedes root hair cell death, it is possible that root hair cells have already died before secondary growth sets in, precluding the detection of dPCD signature genes in this context.

While it remains challenging to image root hair cells in a soil context, new techniques as synchrotron-based X-ray computed microtomography might enable us to follow root hair senescence in their natural context (Duddek et al., 2024). It is tempting to speculate that the reduction of root hairs seen in autophagy mutants would seriously compromise water and mineral acquisition in soil. If modulating root hair life span by targeting autophagy and dPCD processes would decrease the need for fertilizers and increase drought resilience, our results can point out exciting new directions in engineering these desirable trait in crop plants.

## Acknowledgements

We thank Yu Yang (ZEISS Sigma 360 SEM, ZEISS Microscopy Customer Center Shanghai (ZMCCSH)) provided SEM imaging support. This research was financially supported by the Natural Science Foundation of China (32400293 to Q.F.), by the Natural Science Foundation of Shandong Province (2024HWYQ-060 to Q.F.), Tai-Shan Scholar Program of the Shandong Provincial Government (NO.tsqn202312145 to Q.F.), the European Research Council (ERC) StG PROCELLDEATH 639234 and CoG EXECUT.ER 864952 to M.K.N..

## Author contributions

Q.F., M.K.N., and Y.D. analyzed the data, designed the experiments and wrote the article. Q.F. and X.W. prepared the figures. Q.F. and S.Z performed all experiments, except for the following: construction of the vectors *pE7:ATG5-mCherry*, *pIRX1:ATG5-mCherry* and obtaining of the transgenic lines and crosses of *atg* mutants with *sid2*, *coi1*, and *ein2* were done by X.W.; the introducing PCD-reporter line *pRNS3>>HA2-GFP* into *atg* mutants and the quantification of root hair cells expressing of PASPA3 in CRISPR-Cas9 based root hair specific knock out *ATG2* lines were done by Y.L.; mitochondrial leakage experiment was confirmed by J.Z.; M.K.N. performed the imaging of *pANAC046:NLS-tdTOMATO* and *pANAC087:NLS-tdTOMATO* and imaging of *pUBQ10:TOIM* in imaging chamber.

## Materials and methods

### Plant materials, growth, and transformation

*Arabidopsis thaliana* seedlings were grown vertically on 1/2 Murashige and Skoog (MS) medium (2.15 g/L MS salts, 1 g/L sucrose, pH 5.8 [KOH], and 1% (w/v) plant agar) in day/night conditions (16 h light, 8 h dark, 21 °C) before analysis, unless stated otherwise.

The Arabidopsis thaliana *atg2-2* mutant allele (EMS, Gln803stop) was reported by (Wang et al., 2011); *atg5-1* (SAIL_129_B07) was reported by (Thompson et al., 2005); *atg7-2* (GABI_655B06) was reported by (Hofius et al., 2009), *anac046 anac087* was reported by (Huysmans et al., 2018). The Arabidopsis lines *pUBQ10::YFP-ATG8A* and *pUBQ10:TOIM* in wild type and *atg5-1* were reported by (Feng et al., 2022), the *pANAC046:NLS-tdTOMATO* and *pANAC087:NLS-tdTOMATO* were reported by (Huysmans et al., 2018). The *pPASPA3>>H2A-GFP* (Olvera-Carrillo et al., 2015), *pPASPA3:TOIM* (Fendrych et al., 2014), *pRNS3>>H2A-GFP* (Olvera-Carrillo et al., 2015), *pUBQ10:YFP-ATG8A* and *pUBQ10:COX4-GFP* (Wang et al., 2024) were introduced into *atg* mutants by cross. Arabidopsis mutant alleles *sid2-1* (Nawrath and Métraux, 1999), *coi1-2* (Huang et al., 2014), and *ein2-1* (Alonso et al., 1999) were used for intermutant crosses with either *pPASPA3:TOIM/atg2-2* or *atg5-1*. *ATG2* CRISPR lines and *ATG5* complement lines were created by stable Arabidopsis transformation using the floral dipping method described before (Clough and Bent, 1998).

### Cloning

The 726 bp length fragment of pE7 and the 2946 bp length fragment of pRHD6 were amplified by PCR using p524/p525 and p4540/p4541, respectively. These purified PCR fragments were inserted into pGG-A-ccdb-B module vector via a Golden Gate reaction and obtained Golden Gate entry modules pGG-A-pE7-B and pGG-A-pRHD6-B. pGG-B-Linker-C, pGG-C-Cas9-D, pGG-D-P2A-mCherry-NLS-E, pGG-D-P2A-mTagBFP2-NLS-E, pGG-E-G7T-F, pGG-F-pATU6-26-AarI-G, were reported previously (Decaestecker et al., 2019). pGG-C-ANAC046-D and pGG-F-linker-G were obtained from the VIB-UGent plasmid repository (https://gatewayvectors.vib.be). These entry modules were assembled in pFASTRK-AG with FAST selection marker, resulting in the destination vector pFASTRK-pE7-Cas9-P2A-GFP-NLS-pATU6-26-AarI, pFASTRK-pRHD6-Cas9-P2A-GFP-NLS-pATU6-26-AarI and pFASTRK-pE7:ANAC046-P2A-mTagBFP2-NLS. The destination vector pFASTG-pUBI-Cas9 was reported previously (Decaestecker et al., 2019). Fragment gRNA1-pATU6-26-gRNA2 (ATG2 target) was amplified by PCR as described previously (Feng et al., 2022). These purified PCR fragments were inserted into pFASTR-pE7-Cas9-P2A-GFP-NLS-pATU6-26-AarI or pFASTR-pRHD6-Cas9-P2A-GFP-NLS-pATU6-26-AarI or pFASTG-pUBI-Cas9 destination vector via a Golden Gate reaction. The resulting vectors were named *pE7:Cas9;ATG2*, *pRHD6:Cas9;ATG2* and *pUBI:Cas9;ATG2*.

Gateway entry modules L4-pUBQ10-R1 and L1-ATG5-L2 were described previous (Huysmans et al., 2018; Feng et al., 2022). The fragments of mCherry, pE7 and pIRX1 were amplified by PCR using p73/p74, p508/p509 and p510/p507, respectively. The fragment of mCherry was inserted into pDONRP2R-P3 via BP reaction, resulting in entry vector, R2-mCherry-L3. The fragments of pE7 and pIRX1 were inserted into pDONRP4P1R via BP reaction, resulting in entry vector, L4-pE7-R1 and L4-pIRX1-R1. Entry vectors L1-ATG5-L2, R2-mCherry-L3 were assembled with L4-pUBQ10-R1 or L4-pE7-R1 or L4-pIRX1-R1 into pFASTGB-34GW destination vector via LR reaction and resulting in vectors *pFASTGB-pUBQ10:ATG5-mCherry*, *pFASTGB-pE7:ATG5-mCherry*, *pFASTGB-pIRX1:ATG5-mCherry*, respectively.

All primers for cloning are listed in Supplemental Table 1.

### Pharmacological treatments and imaging

For the *pPASPA3>>H2A-GFP* and *pRNS3>>H2A-GFP* confocal imaging, seedlings were mounted on a glass slide in liquid 1/2 MS and imaged on the LSM710 (Zeiss) or LSM880 (Zeiss). For the *pUBQ10:COX4-GFP* confocal imaging, seedling were transferred into imaging chamber 30 min before imaging, and imaged on Nikon AX/AX R. GFP was excited by the 488-nm of the argon laser and detected between 500 and 550 nm.

*pUBQ10:YFP-ATG8A* seedlings were dipped into 1 μM Concanamycin A (ConA) for 8 h, then mounted on a glass side in 1/2 MS medium and imaged on SP8 (Leica). For the imaging of *pUBQ10:YFP-ATG8A* in root hair, seedlings were mounted on a glass slide in 1/2 MS medium and imaged on LSM880. YFP was excited by the 514-nm line of the argon laser and detected between 525 and 580 nm.

For the *pPASPA3:TOIM* confocal imaging, seedlings were mounted on a glass slide in 1/2 MS. For the *pUBQ10:TOIM* root hair imaging, 3-day-old seedlings were transferred into imaging chamber and kept 11 days growing, then imaged on LSM710 (Zeiss). Imaging of ToIM was performed as described before (Fendrych et al., 2014).

*pANAC046:NLS-tdTOMATO* and *pANAC087:NLS-tdTOMATO* were imaged as reported previously (Huysmans et al., 2018).

For the *pRHD6:Cas9;ATG2* and *pE7:Cas9;ATG2* in *pPASPA3>>H2A-GFP*, seedlings were mounted on a glass slide in 1/2 MS medium and imaged on LSM880. GFP and mCherry were detected in the different tracks. GFP was excited by the 488-nm of the argon laser and detected between 500 and 550 nm. mCherry was excited by the 561-nm and detected between 600 and 700 nm.

For root hair morphology, roots were imaged directly from petri dish using OLYMPUS SZX16. Quantification of root hair length and identity was performed as described (Huang et al., 2013). All images were processed and analyzed using Fiji (https://fiji.sc/) (Schindelin et al., 2012).

For quantification of root hair survival ratio, 14-day-old primary roots were marked every 1 cm from the junction of hypocotyl and root to root tip. In the *atg* mutants, the area close to root tip in which root hairs start to undergo collapse was marked as stage 1, the lower area was labeled as minus 1. The corresponding position in WT was labeled the same. For each stage, root hairs were imaged using OLYMPUS SZX16. The number of normal and collapsed root hairs was counted using Fiji. The survival ratio was obtained by the number of normal root hair divided the number of total root hair.

### RNA extraction and digital-droplet PCR (ddPCR)

Total RNAs were isolated using a Qiagen RNeasy plant mini kit according to the manufacturer’s instructions. About 20-30 roots from 6-day-old seedlings were harvested for RNA extraction per each genotype. ddPCR was performed using the QIAcuity EG PCR Kit according to the manufacturer’s instructions. Primers for PCD-associated genes were reported previously (Olvera-Carrillo et al., 2015; Huysmans et al., 2018). GAPDH was used as internal controls (Ryu et al., 2010). RNA extractions and ddPCR were performed for three biological replicates for each genotype, and two or three technical replicates were performed for each sample. The data were normalized against wild-type. All primers for ddPCR are listed in Supplemental Table 1.

### Quantification and Statistical Analysis

The statistical details of experiments can be found in the corresponding figure legends. The results of statistical tests can be found in the corresponding Results section. Statistical tests were carried out using GraphPad Prism 9.0.0.

## Supplemental data

**Supplemental Figure 1.**
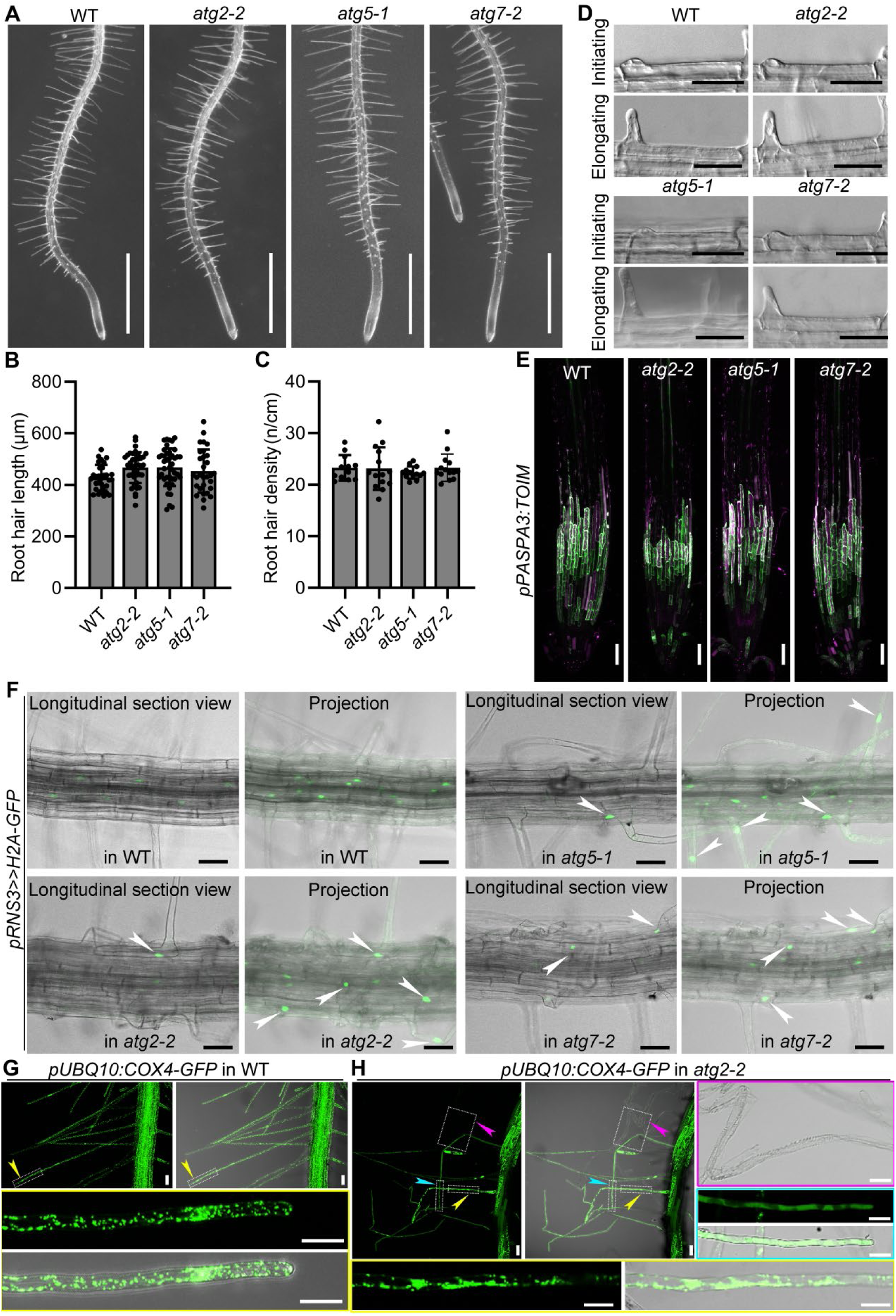
Root hair development and marker gene expression in the wild-type (WT) and *atg* mutants. A, Representative primary roots from 5-day-old seedling: WT, *atg2-2*, *atg5-1* and *atg7-2* seedlings. B-C, The quantification of root hair length (B) or intensity (C) of WT and *atg* mutants. Results shown are means ± standard deviation (SD). Three independent experiments involving 30-42 roots were performed (one-way ANOVA, Tukey’s multiple comparisons test, P > 0.05). D, Differential interference contrast images of initiating (upper panel) or elongating (lower panel) root hairs from WT and *atg* mutants. E, The *pPASPA3:TOIM* marker showed expression and cell death in the distal and proximal root cap cells and xylem of the *atg* mutants, similar to WT. The eGFP signal is shown in green in the cytoplasm, whereas the mRFP signal is shown in magenta in the vacuole. Merge of both signals indicates vacuolar collapse, a hallmark of cell death. F, Representative confocal images of *pRNS3>>H2A-GFP* signals in 5-day-old WT, *atg2-2*, *atg5-1* and *atg7-2*. White arrowheads indicate that H2A-GFP signals were observed in root hair cells of *atg* mutants. G, Representative confocal images of mitochondria matrix marker *pUBQ10:COX4-GFP* signals in 6-day-old seedlings from WT and *atg2-2*. The zoomed images are shown of a part region of outlined in white dotted lines. Different colored arrowheads indicate different types of root hairs of *atg2-2* mutants. Magenta arrowheads indicate the dead root hairs. Cyan arrowhead indicate the dying root hairs. Yellow arrowheads indicate the living root hairs. Images are shown in z-projection, except the zoomed images in different color lined boxes. Bars = 100 mm for A, 50 μm for D, E, F and G, 20 μm for the zoomed images in G. Related to Figure 2.

**Supplemental Figure 2.**
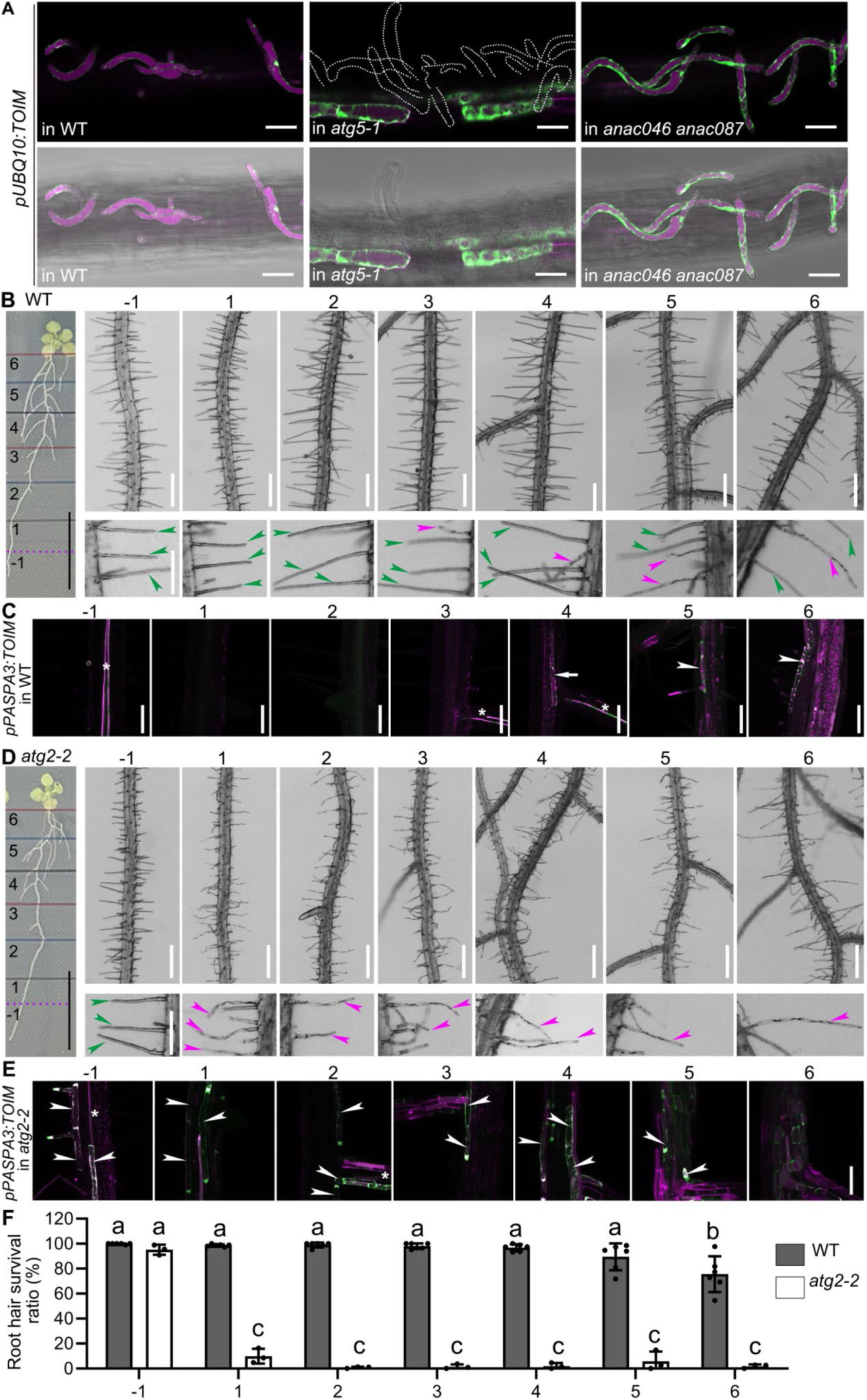
a*t*g mutants show earlier root hair cell death than the wild type. A, *pUBQ10:TOIM* expression in WT, *atg5-1* and *anac046 anac087* was imaged in the growing chamber. Root hair cells close to first emerged lateral root in 14-day-old seedlings were imaged. *atg5-1* mutant shows premature root hair cell death compared with WT. EGFP signal is shown in green in the cytoplasm, and mRFP signal is shown in magenta in the vacuole. Bars = 50 μm. B and D, Illustration of a 14-day-old seedling showing the positions corresponding to the different stages (left panel of B and D). Roots at different positions of the root corresponding to from stages -1 to 6 are shown on the right. Lower panel shows a zoomed image of a part region of the upper panel. Green arrowheads indicate living root hair, while magenta arrowheads indicate the dead root hair. Bars = 1.5 cm for left panel, 500 μm for upper panel and 250 μm for lower panel. C and E, Representative confocal images of roots from 14-day-old WT (C) and *atg* mutants (E) expressing *pPASPA3:TOIM*. White arrowheads indicate that TOIM signals were observed in root hair cells. White arrows indicate that TOIM signals were observed in cortex cells suggesting the periderm growth. White asterisks indicate TOIM signals were observed in xylem cells. Bars = 100 μm. F, The quantification of root hair survival ratio. Results shown are means ± SD. At least three independent experiments involving 400-550 root hairs were performed. Means with different letters are significantly different (two-way ANOVA, Tukey’s multiple comparisons test, P < 0.05). Related to Figure 2.

**Supplemental Figure 3.**
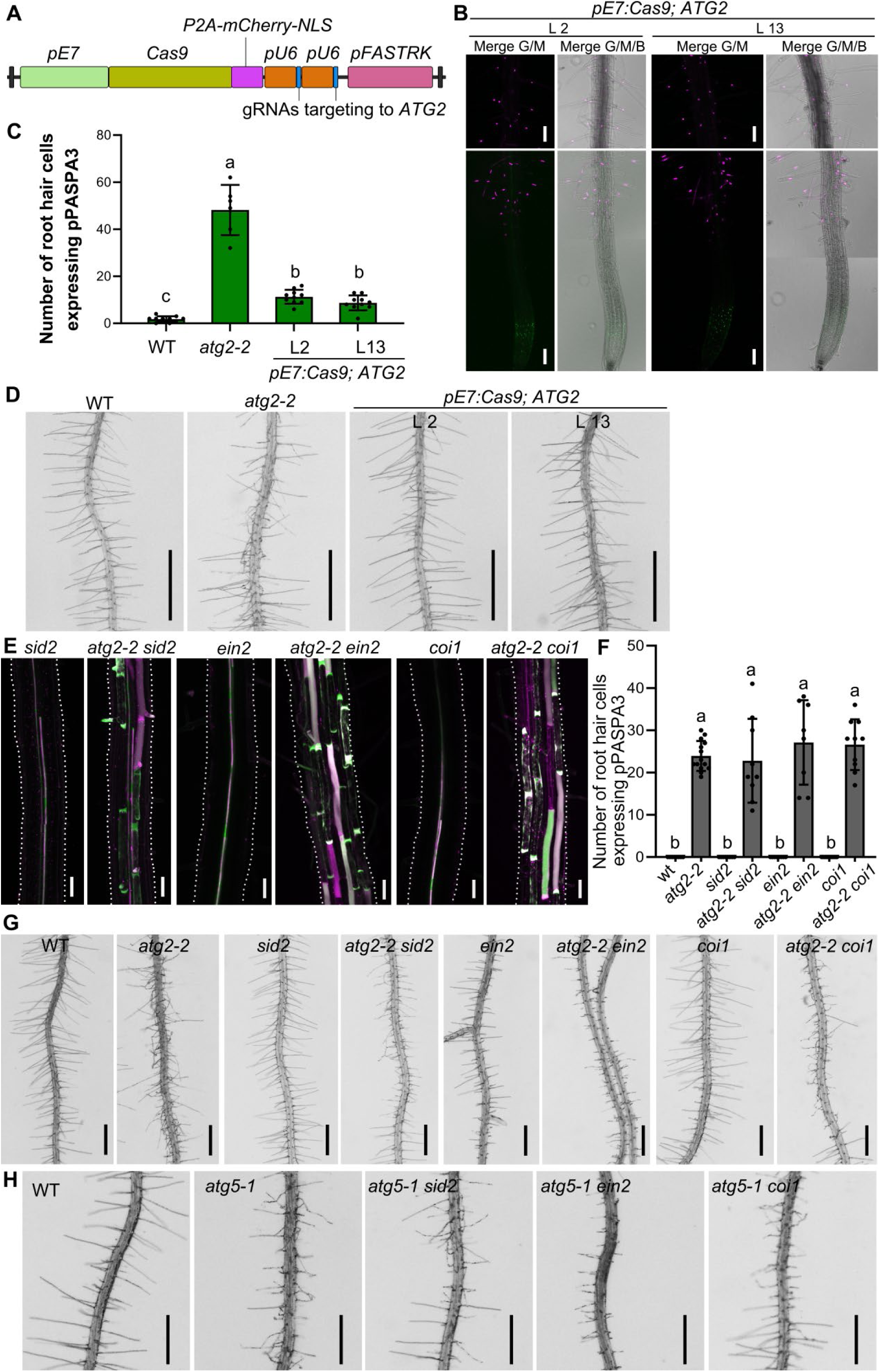
Root hair-specific knock out *ATG2*, and independency of the premature root hair phenotype in *atg2* and *atg5* from SA, JA and ethylene signaling. A, Schematic representation of the vector: *pE7:Cas9;ATG2.* The Cas9-P2A-mCherry is driven by root hair specific promoter pE7, and two gRNAs targeting to *ATG2* are inserted. B, Representative confocal images of two independent lines of *pE7:Cas9;ATG2* in *pPASPA3>>H2A-GFP* background. C, The quantification of root hair cell expressing *PASPA3*. Results shown are means ± SD. In total, 6-10 roots were analyzed for each genotype. Means with different letters are significantly different (one-way ANOVA, Tukey’s multiple comparisons test, P < 0.05). D, Representative primary roots from 7-day-old seedlings: WT, *atg2-2* and two independent lines of *pE7:Cas9; ATG2*. E, Representative confocal images of roots from 5-day-old seedlings expressing *pPASPA3:TOIM*: *sid2*, *atg2-2 sid2, ein2*, *atg2-2 ein2*, *coi1* and *atg2-2 coi1*. White dotted lines point the profile of root. F, The quantification of root hair cell expressing *PASPA3*. Results shown are means ± SD. In total, 8-14 roots were analyzed for each genotype. Means with different letters are significantly different (one-way ANOVA, Tukey’s multiple comparisons test, P < 0.05). G, Representative primary roots from 7-day-old seedlings: WT, *atg2-2*, *sid2*, *atg2-2 sid2, ein2*, *atg2-2 ein2*, *coi1* and *atg2-2 coi1*. H, Representative primary roots from 7-day-old seedlings: WT, *atg5-1*, *atg5-1 sid2*, *atg5-1 ein2* and *atg5-1 coi1*. Bars = 100 μm for B, 1 mm for D, 50 μm for E, 500 μm for G and H. Related to Figure 3.

**Supplemental Figure 4.**
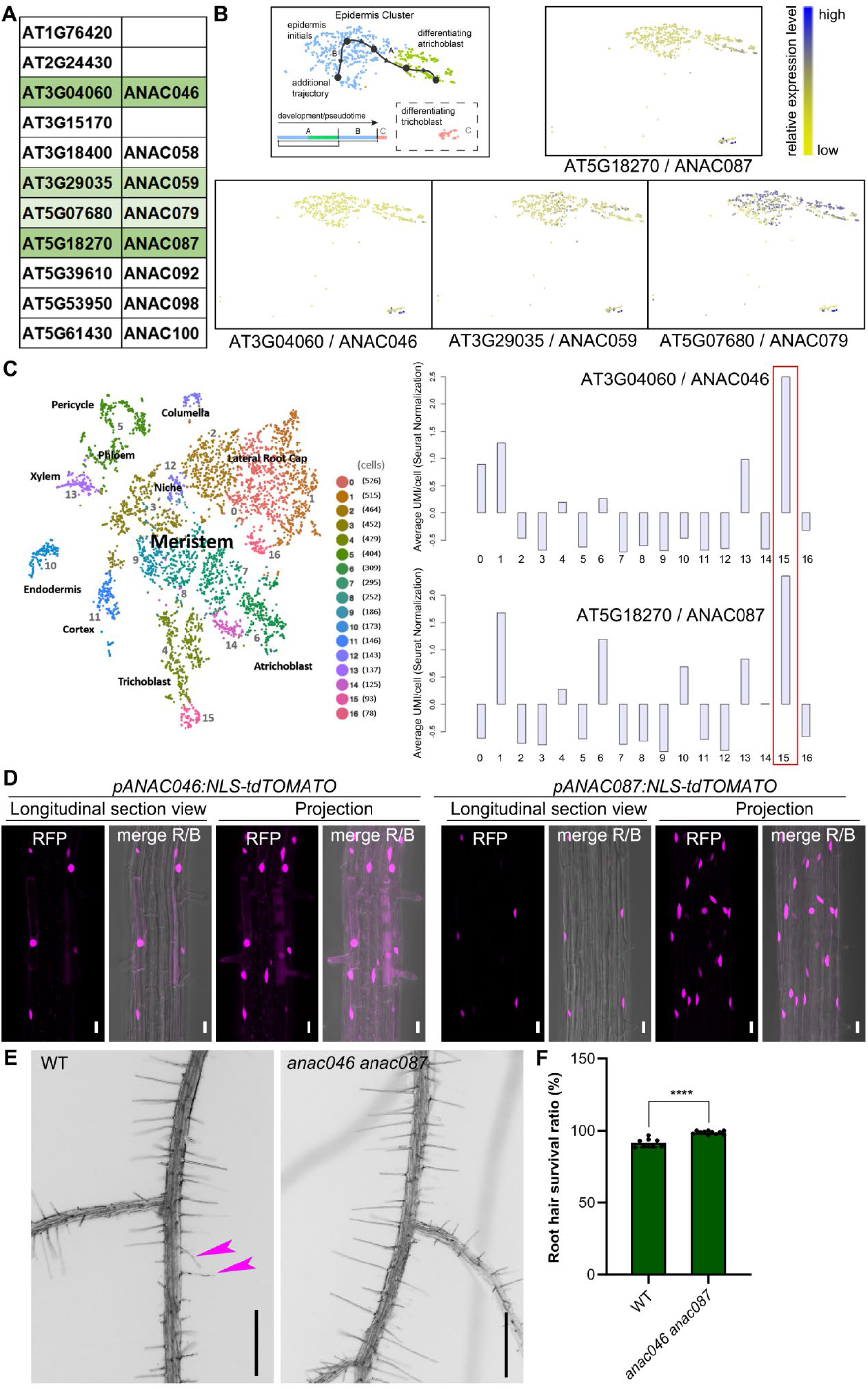
Expression analysis of *ANAC046* and *ANAC087* and root hair longevity analysis of *anac046 anac087* mutants. A, ANAC046 family members via PLAZA blast. Green color high light indicate which is detected in sc-RNAseq data, shown as in B. B, Expression pattern of *ANAC046*, *ANAC059*, *ANAC079* and *ANAC087* in epidermis cluster in published sc-RNAseq dataset (<u>Plant sc-Atlas</u> <u>(ugent.be)</u>). C, Expression pattern of *ANAC046* and *ANAC087* in published sc-RNAseq dataset (<u>zmbp-</u> <u>resources.uni-tuebingen.de/timmermans/plant-single-cell-browser/</u>). Red box indicates root hair cluster. D, Representative confocal images of roots from 5-day-old seedlings expressing *pANAC046:NLS-tdTOMATO and pANAC087:NLS-tdTOMATO.* Merges of the RFP/bright field (R/B) are shown at the right. TdTOMATO is shown in magenta. Bars = 20 μm. E, Representative primary roots from 14-day-old seedlings close to hypocotyl: WT and *anac046 anac087*. Magenta arrowheads indicate the collapsed root hair. Bars = 500 μm. F, The quantification of root hair survival ratio. Results shown are means ± SD. In total, 4000-5300 root hairs from at least 8 roots were analyzed for each genotype. **** indicates a significant difference (t test, P < 0.0001). Related to Figure 4.

**Supplemental Figure 5.**
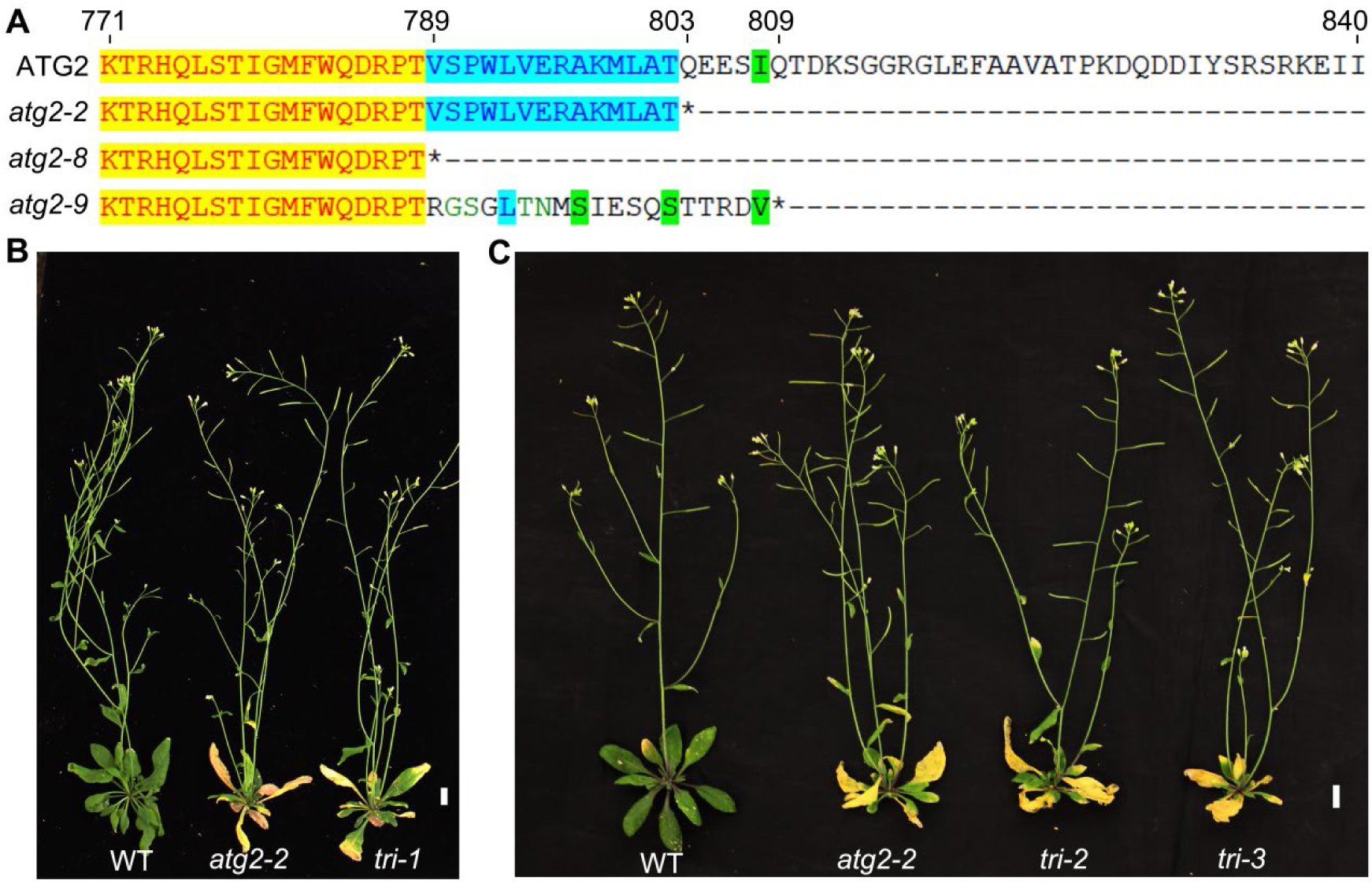
Amino acid sequence alignment of *atg2* mutants, and leaf senescence analysis of *atg2-2* versus triple mutants of *atg2 anac046 anac087.* A, Partial amino acid (aa) sequence of ATG2 and the mutants. *atg2-2* has early stop codon at 803 aa. *atg2-8* has early stop codon at 789 aa. While *atg2-9* has different aa from 789 aa on and has early stop codon at 809 aa. B-C, Representative images from 5-week-old plants: WT, *atg2-2*, *tri-1* (*atg2-2 anac046 anac087*), *tri-2* (*atg2-8 anac046 anac087*), *tri-3* (*atg2-9 anac046 anac087*). Bars = 1 cm. Related to Figure 4.

**Supplemental table 1.**
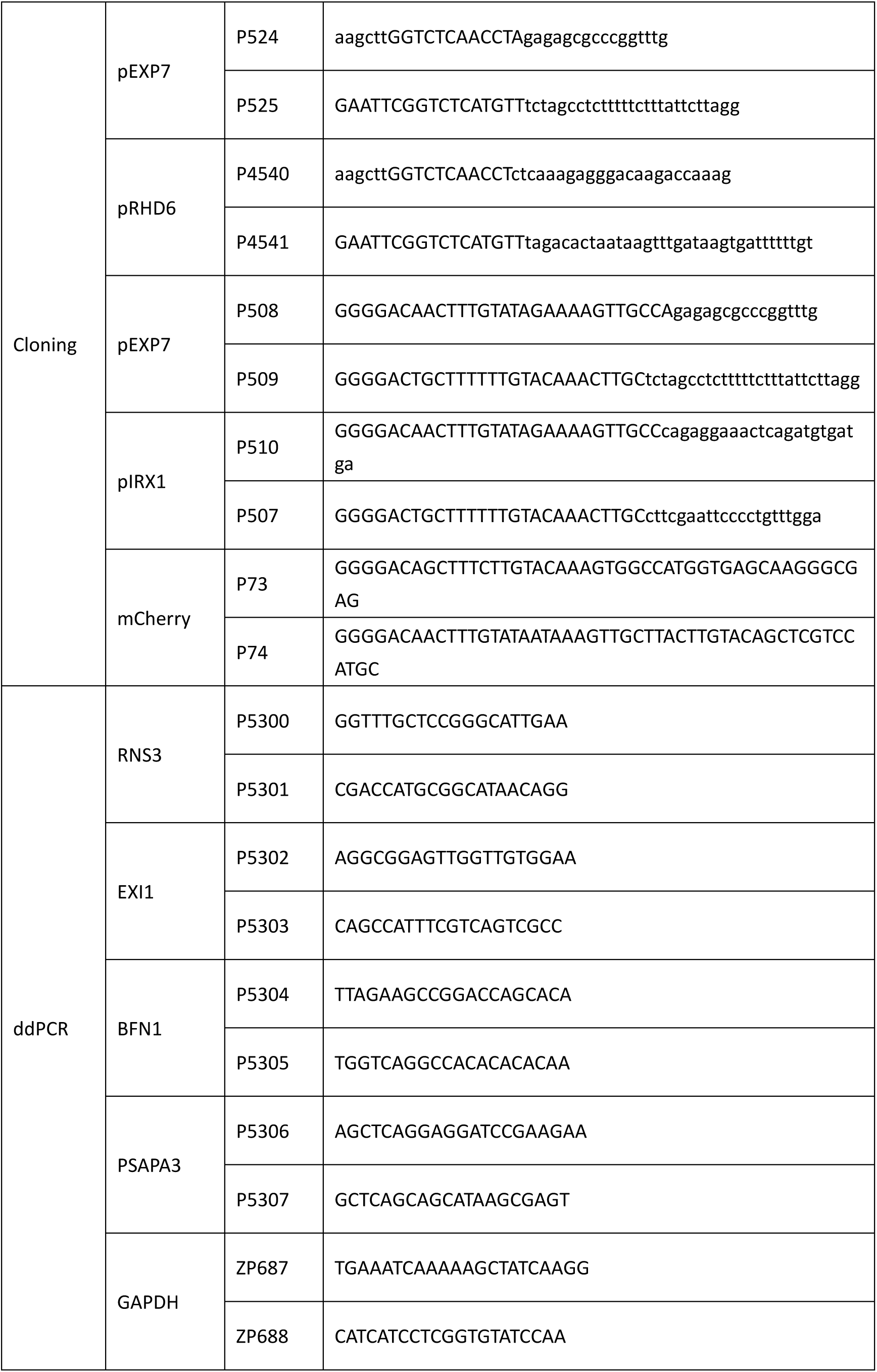
Primers were used in this paper.

## References

Agbemafle W, Wong MM, and Bassham DC. Transcriptional and post-translational regulation of plant autophagy. J Exp Bot. 2023:74(19): 6006–6022.

Alonso JM, Hirayama T, Roman G, Nourizadeh S, and Ecker JR. EIN2, a bifunctional transducer of ethylene and stress responses in Arabidopsis. Science. 1999:284(5423): 2148–2152.

Bates TR, and Lynch JP. Stimulation of root hair elongation in Arabidopsis thaliana by low phosphorus availability. 1996:19(5): 529–538.

Bollier N, Andrade Buono R, Jacobs TB, and Nowack MK. Efficient simultaneous mutagenesis of multiple genes in specific plant tissues by multiplex CRISPR. Plant Biotechnol J. 2021:19(4): 651–653.

Cai G, and Ahmed MA. The role of root hairs in water uptake: recent advances and future perspectives. J Exp Bot. 2022:73(11): 3330–3338.

Cassidy LD, and Narita M. Autophagy at the intersection of aging, senescence, and cancer. Mol Oncol. 2022:16(18): 3259–3275.

Cho HT, and Cosgrove DJ. Regulation of root hair initiation and expansin gene expression in Arabidopsis. Plant Cell. 2002:14(12): 3237–3253.

Choi HS, and Cho HT. Root hairs enhance Arabidopsis seedling survival upon soil disruption. Sci Rep. 2019:9(1): 11181.

Clough SJ, and Bent AF. Floral dip: a simplified method for Agrobacterium-mediated transformation of Arabidopsis thaliana. Plant J. 1998:16(6): 735–743.

Courtois-Moreau CL, Pesquet E, Sjödin A, Muñiz L, Bollhöner B, Kaneda M, Samuels L, Jansson S, and Tuominen H. A unique program for cell death in xylem fibers of Populus stem. Plant J. 2009:58(2): 260–274.

Cui S, Suzaki T, Tominaga-Wada R, and Yoshida S. Regulation and functional diversification of root hairs. Semin Cell Dev Biol. 2018:83: 115–122.

Daneva A, Gao Z, Van Durme M, and Nowack MK. Functions and Regulation of Programmed Cell Death in Plant Development. Annu Rev Cell Dev Biol. 2016:32: 441–468.

Decaestecker W, Buono RA, Pfeiffer ML, Vangheluwe N, Jourquin J, Karimi M, Van Isterdael G, Beeckman T, Nowack MK, and Jacobs TB. CRISPR-TSKO: A Technique for Efficient Mutagenesis in Specific Cell Types, Tissues, or Organs in Arabidopsis. Plant Cell. 2019:31(12): 2868–2887.

Denyer T, Ma X, Klesen S, Scacchi E, Nieselt K, and Timmermans MCP. Spatiotemporal Developmental Trajectories in the Arabidopsis Root Revealed Using High-Throughput Single-Cell RNA Sequencing. Dev Cell. 2019:48(6): 840–852.e845.

Doll NM, Van Hautegem T, Schilling N, De Rycke R, De Winter F, Fendrych M, and Nowack MK. Endosperm cell death promoted by NAC transcription factors facilitates embryo invasion in Arabidopsis. Curr Biol. 2023:33(17): 3785–3795 e3786.

Duddek P, Papritz A, Ahmed MA, Lovric G, and Carminati A. Observations of root hair patterning in soils: Insights from synchrotron-based X-ray computed microtomography. Plant and Soil. 2024:503(1): 331–348.

Duddek P, Ahmed MA, Javaux M, Vanderborght J, Lovric G, King A, and Carminati A. The effect of root hairs on root water uptake is determined by root-soil contact and root hair shrinkage. New Phytol. 2023:240(6): 2484–2497.

Duddek P, Carminati A, Koebernick N, Ohmann L, Lovric G, Delzon S, Rodriguez-Dominguez CM, King A, and Ahmed MA. The impact of drought-induced root and root hair shrinkage on root-soil contact. Plant Physiol. 2022:189(3): 1232–1236.

Escamez S, Andre D, Zhang B, Bollhoner B, Pesquet E, and Tuominen H. METACASPASE9 modulates autophagy to confine cell death to the target cells during Arabidopsis vascular xylem differentiation. Biol Open. 2016:5(2): 122–129.

Escamez S, Stael S, Vainonen JP, Willems P, Jin H, Kimura S, Van Breusegem F, Gevaert K, Wrzaczek M, and Tuominen H. Extracellular peptide Kratos restricts cell death during vascular development and stress in Arabidopsis. J Exp Bot. 2019:70(7): 2199–2210.

Fendrych M, Van Hautegem T, Van Durme M, Olvera-Carrillo Y, Huysmans M, Karimi M, Lippens S, Guerin CJ, Krebs M, Schumacher K, et al. Programmed cell death controlled by ANAC033/SOMBRERO determines root cap organ size in Arabidopsis. Curr Biol. 2014:24(9): 931–940.

Feng Q, De Rycke R, Dagdas Y, and Nowack MK. Autophagy promotes programmed cell death and corpse clearance in specific cell types of the Arabidopsis root cap. Curr Biol. 2022:32(9): 2110–2119.

Galluzzi L, Vitale I, Aaronson SA, Abrams JM, Adam D, Agostinis P, Alnemri ES, Altucci L, Amelio I, Andrews DW, et al. Molecular mechanisms of cell death: recommendations of the Nomenclature Committee on Cell Death 2018. Cell Death Differ. 2018:25(3): 486–541.

Gao Z, Daneva A, Salanenka Y, Van Durme M, Huysmans M, Lin Z, De Winter F, Vanneste S, Karimi M, Van de Velde J, et al. KIRA1 and ORESARA1 terminate flower receptivity by promoting cell death in the stigma of Arabidopsis. Nat Plants. 2018:4(6): 365–375.

Gilroy S, and Jones DL. Through form to function: root hair development and nutrient uptake. Trends Plant Sci. 2000:5(2): 56–60.

Goh T, Sakamoto K, Wang P, Kozono S, Ueno K, Miyashima S, Toyokura K, Fukaki H, Kang BH, and Nakajima K. Autophagy promotes organelle clearance and organized cell separation of living root cap cells in Arabidopsis thaliana. Development. 2022:149(11).

Grierson C, Nielsen E, Ketelaarc T, and Schiefelbein J. Root hairs. Arabidopsis Book. 2014:12: e0172.

Gross AS, Raffeiner M, Zeng Y, Üstün S, and Dagdas Y. Autophagy in Plant Health and Disease. Annu Rev Plant Biol. 2025.

Hofius D, Schultz-Larsen T, Joensen J, Tsitsigiannis DI, Petersen NH, Mattsson O, Jorgensen LB, Jones JD, Mundy J, and Petersen M. Autophagic components contribute to hypersensitive cell death in Arabidopsis. Cell. 2009:137(4): 773–783.

Hogg BV, Kacprzyk J, Molony EM, O’Reilly C, Gallagher TF, Gallois P, and McCabe PF. An in vivo root hair assay for determining rates of apoptotic-like programmed cell death in plants. Plant Methods. 2011:7(1): 45.

Honkanen S, and Dolan L. Growth regulation in tip-growing cells that develop on the epidermis. Curr Opin Plant Biol. 2016:34: 77–83.

Huang H, Wang C, Tian H, Sun Y, Xie D, and Song S. Amino acid substitutions of GLY98, LEU245 and GLU543 in COI1 distinctively affect jasmonate-regulated male fertility in Arabidopsis. Sci China Life Sci. 2014:57(1): 145–154.

Huysmans M, Buono RA, Skorzinski N, Radio MC, De Winter F, Parizot B, Mertens J, Karimi M, Fendrych M, and Nowack MK. NAC Transcription Factors ANAC087 and ANAC046 Control Distinct Aspects of Programmed Cell Death in the Arabidopsis Columella and Lateral Root Cap. Plant Cell. 2018:30(9): 2197–2213.

Kacprzyk J, Burke R, Armengot L, Coppola M, Tattrie SB, Vahldick H, Bassham DC, Bosch M, Brereton NJB, Cacas JL, et al. Roadmap for the next decade of plant programmed cell death research. New Phytol. 2024:242(5): 1865–1875.

Keyes SD, Zygalakis KC, and Roose T. An Explicit Structural Model of Root Hair and Soil Interactions Parameterised by Synchrotron X-ray Computed Tomography. Bull Math Biol. 2017:79(12): 2785–2813.

Kim HJ, Hong SH, Kim YW, Lee IH, Jun JH, Phee BK, Rupak T, Jeong H, Lee Y, Hong BS, et al. Gene regulatory cascade of senescence-associated NAC transcription factors activated by ETHYLENE-INSENSITIVE2-mediated leaf senescence signalling in Arabidopsis. J Exp Bot. 2014:65(14): 4023–4036.

Kubo M, Udagawa M, Nishikubo N, Horiguchi G, Yamaguchi M, Ito J, Mimura T, Fukuda H, and Demura T. Transcription switches for protoxylem and metaxylem vessel formation. Genes Dev. 2005:19(16): 1855–1860.

Kwon SI, Cho HJ, Jung JH, Yoshimoto K, Shirasu K, and Park OK. The Rab GTPase RabG3b functions in autophagy and contributes to tracheary element differentiation in Arabidopsis. Plant J. 2010:64(1): 151–164.

Li E, Zhang YL, Qin Z, Xu M, Qiao Q, Li S, Li SW, and Zhang Y. Signaling network controlling ROP-mediated tip growth in Arabidopsis and beyond. Plant Commun. 2023:4(1): 100451.

Li L, Tan K, Tang XG, Chao XT, Wen CX, D. BZ, Feng HL, Liu WZ, and Su H. Characterization of Programmed Cell Death During the Senescence of Root Hairs in Arabidopsis. 2016:51(2): 194–201.

Liao CY, Wang P, Yin Y, and Bassham DC. Interactions between autophagy and phytohormone signaling pathways in plants. FEBS Lett. 2022:596(17): 2198–2214.

Liu Y, Schiff M, Czymmek K, Talloczy Z, Levine B, and Dinesh-Kumar SP. Autophagy regulates programmed cell death during the plant innate immune response. Cell. 2005:121(4): 567–577.

Marshall RS, and Vierstra RD. Autophagy: The Master of Bulk and Selective Recycling. Annu Rev Plant Biol. 2018:69: 173–208.

Masclaux-Daubresse C, Chen Q, and Havé M. Regulation of nutrient recycling via autophagy. Curr Opin Plant Biol. 2017:39: 8–17.

Menand B, Yi K, Jouannic S, Hoffmann L, Ryan E, Linstead P, Schaefer DG, and Dolan L. An ancient mechanism controls the development of cells with a rooting function in land plants. Science. 2007:316(5830): 1477–1480.

Nawrath C, and Métraux JP. Salicylic acid induction-deficient mutants of Arabidopsis express PR-2 and PR-5 and accumulate high levels of camalexin after pathogen inoculation. Plant Cell. 1999:11(8): 1393–1404.

Oda-Yamamizo C, Mitsuda N, Sakamoto S, Ogawa D, Ohme-Takagi M, and Ohmiya A. The NAC transcription factor ANAC046 is a positive regulator of chlorophyll degradation and senescence in Arabidopsis leaves. Sci Rep. 2016:6: 23609.

Ohashi-Ito K, Oda Y, and Fukuda H. Arabidopsis VASCULAR-RELATED NAC-DOMAIN6 directly regulates the genes that govern programmed cell death and secondary wall formation during xylem differentiation. Plant Cell. 2010:22(10): 3461–3473.

Olvera-Carrillo Y, Van Bel M, Van Hautegem T, Fendrych M, Huysmans M, Simaskova M, van Durme M, Buscaill P, Rivas S, Coll NS, et al. A Conserved Core of Programmed Cell Death Indicator Genes Discriminates Developmentally and Environmentally Induced Programmed Cell Death in Plants. Plant Physiol. 2015:169(4): 2684–2699.

Ryu MY, Cho SK, and Kim WT. The Arabidopsis C3H2C3-type RING E3 ubiquitin ligase AtAIRP1 is a positive regulator of an abscisic acid-dependent response to drought stress. Plant Physiol. 2010:154(4): 1983–1997.

Schindelin J, Arganda-Carreras I, Frise E, Kaynig V, Longair M, Pietzsch T, Preibisch S, Rueden C, Saalfeld S, Schmid B, et al. Fiji: an open-source platform for biological-image analysis. Nat Methods. 2012:9(7): 676–682.

Shishkova S, and Dubrovsky JG. Developmental programmed cell death in primary roots of Sonoran Desert Cactaceae. Am J Bot. 2005:92(9): 1590–1594.

Tan K, Wen C, Feng H, Chao X, and Su H. Nuclear dynamics and programmed cell death in Arabidopsis root hairs. Plant Sci. 2016:253: 77–85.

Taylor NG, Laurie S, and Turner SR. Multiple cellulose synthase catalytic subunits are required for cellulose synthesis in Arabidopsis. Plant Cell. 2000:12(12): 2529–2540.

Teper-Bamnolker P, Danieli R, Peled-Zehavi H, Belausov E, Abu-Abied M, Avin-Wittenberg T, Sadot E, and Eshel D. Vacuolar processing enzyme translocates to the vacuole through the autophagy pathway to induce programmed cell death. Autophagy. 2021:17(10): 3109–3123.

Thompson AR, Doelling JH, Suttangkakul A, and Vierstra RD. Autophagic nutrient recycling in Arabidopsis directed by the ATG8 and ATG12 conjugation pathways. Plant Physiol. 2005:138(4): 2097–2110.

Tunc CE, and von Wirén N. Hidden aging: the secret role of root senescence. Trends in plant science. 2025.

Turner SR, and Somerville CR. Collapsed xylem phenotype of Arabidopsis identifies mutants deficient in cellulose deposition in the secondary cell wall. Plant Cell. 1997:9(5): 689–701.

Üstün S, Hafrén A, and Hofius D. Autophagy as a mediator of life and death in plants. Curr Opin Plant Biol. 2017:40: 122–130.

Vargas-Hernández BY, Núñez-Muñoz L, Calderón-Pérez B, Xoconostle-Cázares B, and Ruiz-Medrano R. The NAC Transcription Factor ANAC087 Induces Aerial Rosette Development and Leaf Senescence in Arabidopsis. Front Plant Sci. 2022:13: 818107.

Vissenberg K, Claeijs N, Balcerowicz D, and Schoenaers S. Hormonal regulation of root hair growth and responses to the environment in Arabidopsis. J Exp Bot. 2020:71(8): 2412–2427.

Wang J, Bollier N, Buono RA, Vahldick H, Lin Z, Feng Q, Hudecek R, Jiang Q, Mylle E, Van Damme D, et al. A developmentally controlled cellular decompartmentalization process executes programmed cell death in the Arabidopsis root cap. Plant Cell. 2024:36(4): 941–962.

Wang Y, Nishimura MT, Zhao T, and Tang D. ATG2, an autophagy-related protein, negatively affects powdery mildew resistance and mildew-induced cell death in Arabidopsis. Plant J. 2011:68(1): 74–87.

Wendrich JR, Yang B, Vandamme N, Verstaen K, Smet W, Van de Velde C, Minne M, Wybouw B, Mor E, Arents HE, et al. Vascular transcription factors guide plant epidermal responses to limiting phosphate conditions. Science. 2020:370(6518).

Won SK, Lee YJ, Lee HY, Heo YK, Cho M, and Cho HT. Cis-element- and transcriptome-based screening of root hair-specific genes and their functional characterization in Arabidopsis. Plant Physiol. 2009:150(3): 1459–1473.

Woo HR, Kim HJ, Lim PO, and Nam HG. Leaf Senescence: Systems and Dynamics Aspects. Annu Rev Plant Biol. 2019:70: 347–376.

Wunderling A, Ripper D, Barra-Jimenez A, Mahn S, Sajak K, Targem MB, and Ragni L. A molecular framework to study periderm formation in Arabidopsis. New Phytol. 2018:219(1): 216–229.

Xiao S, Liu L, Zhang Y, Sun H, Zhang K, Bai Z, Dong H, and Li C. Fine root and root hair morphology of cotton under drought stress revealed with RhizoPot. 2020:206(6): 679–693.

Yoshimoto K, Jikumaru Y, Kamiya Y, Kusano M, Consonni C, Panstruga R, Ohsumi Y, and Shirasu K. Autophagy negatively regulates cell death by controlling NPR1-dependent salicylic acid signaling during senescence and the innate immune response in Arabidopsis. Plant Cell. 2009:21(9): 2914–2927.

Zhang X, Bian A, Li T, Ren L, Li L, Su Y, and Zhang Q. ROS and calcium oscillations are required for polarized root hair growth. Plant Signal Behav. 2022:17(1): 2106410.

